# Interactive effect matters: a combination of herbivory degree and the ratio of generalist to specialist better predicts evolution of plant defense

**DOI:** 10.1101/2022.01.13.476260

**Authors:** Yuanfei Pan, Xiaoyun Pan, Lucas Del Bianco Faria, Bo Li

## Abstract

Herbivory degree and the ratio of generalist to specialist herbivores have long been treated as two important but independent factors in shaping the evolution of plant defense. However, this assumption of independency is poorly supported and has resulted in great controversy in explaining the patterns of plant defense. Here we investigated the possible interaction between herbivory degree and generalist-to-specialist ratio using a cost-benefit model of defense evolution in plants. Our results showed that, with increasing generalist herbivore proportion, plant defense investment increases when herbivory degree is low and decreases when herbivory degree is high. These results provide the first theoretical support for the interactive effect of herbivory degree and ratio of generalist/specialist affecting plant defense, which integrate many of the previous results (e.g. latitudinal patterns of plant defense and defense evolution of invasive plants) and put them into a more general theoretical context.

## Introduction

Plants have evolved a variety of strategies to defend against herbivores. These strategies include secondary metabolites (such as phenols, flavonoids, terpenes and alkaloids), as well as physical traits (such as spines, thorns, and trichomes) (Fritz & Simms 1992). Since herbivore damage may lead to significant lose in plant fitness, it is expected that natural selection should favor high levels of defense (Marquis, 1984, 1992). However, numerous studies have found the existence of intermediate levels of plant defense with substantial variations both among and within plant species (Rausher & Simms, 1989; Vrieling *et al.,* 1993; Mauricio & Rausher, 1997). These patterns support the idea that selection from herbivores on plant defense may vary in both direction and strength and thus there could be trade-offs in shaping the evolution of plant defenses. Studies that tried to explain interspecific or intraspecific variation in plant defense largely fall into two categories.

On one hand, the optimal defense hypothesis (ODH, McKey 1974, 1979; Rhoades 1979; Stamp 2003) explains these patterns with an elegant cost-benefit analysis. It assumes that plant allocates to defense in portion to the herbivory degree it is suffering and plant defense is costly, thus an intermediate level of defense will be favored. Many prominent hypotheses have incorporated the idea from the ODH. For example, the evolution of increased competitive ability hypothesis (EICA, Blossey & Nötzold 1995) predicts that invasive plants should evolve a low level of defense as a result of being released from natural enemies in their introduced range. The latitudinal herbivory-defense hypothesis (LHDH, Coley & Aide 1991) posits that herbivory degree and plant defenses increase toward lower latitudes. A substantial bodies of recent studies testing EICA or LHDH have provided contradictory evidences (EICA: Handley *et al.* 2008; Chun *et al.* 2010; Felker-Quinn *et al.* 2013; LHDH: Moles *et al.* 2011a, b). This is indicating that besides herbivory degree there should be other factors that influence biogeographic patterns of plant defense.

On the other hand, many studies have suggested that generalist (feed on many different plant species) and specialist herbivores (feed on a restricted set of related plant species) may exert opposing selections on plant defense. Typically generalist herbivores are effectively deterred by high concentrations of defense chemicals (Cornell & Hawkins, 2003; Ali & Agrawal, 2012), while some specialist herbivores have evolved to utilize plant chemicals as oviposition cues or feeding stimulants (Macel & Vrieling, 2003; Nieminen *et al.,* 2003). Several specialists even sequester such chemicals and use them for their own defense against natural enemies (Lankau, 2007; Ali & Agrawal, 2012; Züst *et al*., 2018). This contrasting effect of generalist and specialist on plant defense has attracted some theoretical attention over the last century (van der Meijden, 1996) and a few empirical studies tested some of the predictions made by these theoretical analyses (Lankau, 2007; Liu *et al.,* 2018; Zhang *et al.,* 2018). Thus, besides herbivory degree, variation in herbivore composition (e.g. the ratio of specialist to generalist herbivores) through space and time could be another important factor for maintenance of genetic variation in defensive traits. In some extreme cases, if specialist herbivores are absent, as is the case with some introduced plant species in their introduced range, selection may lead to rapid increases in the levels of chemical defense (shifting defense hypothesis, SDH, Müller-Schärer *et al.* 2004; Joshi & Vrieling 2005; Zhang *et al.* 2018).

In summary, herbivory degree (e.g. the total leaf herbivory) and the ratio of generalist to specialist have long been treated as two important but independent factors in shaping the evolution of plant defense. Reasons for such a knowledge gap could be twofold. Theoretically, the graphical model by van der Meijden (1996) simply assumed that herbivory degree to a non-defended plant genotype is fixed, so that this theoretical study could not explore such interaction. And technically, in order to detect such interaction with experiments, a response-surface design with treatment gradient of both herbivore degree and the ratio of generalist to specialist is needed (e.g. Fig. 1a). But conducting such experiments is labor-intensive.

**Figure 1.**
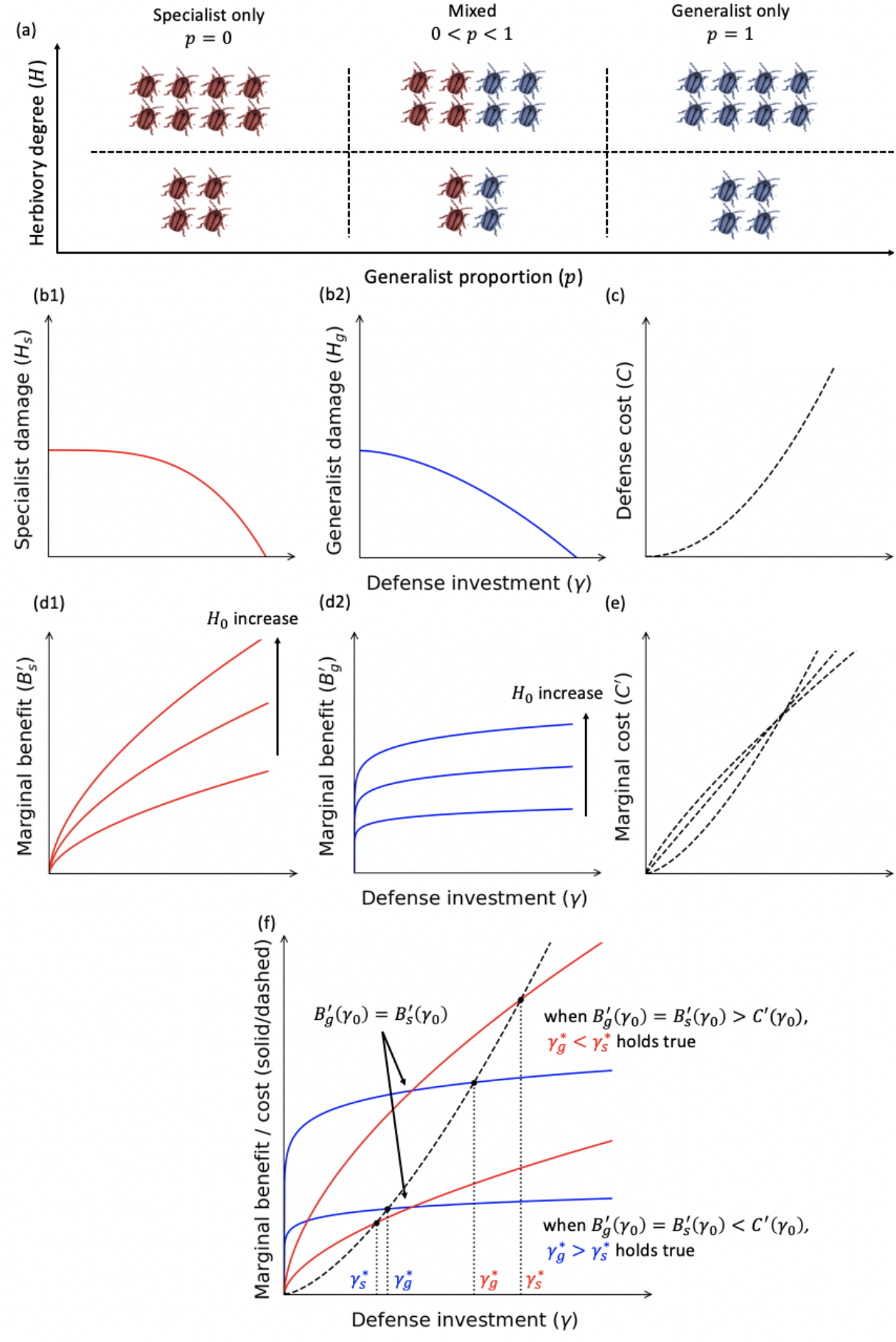
Conceptual framework for explaining the possible interactive effect between herbivory degree (*H*) and the ratio of generalist to specialist (*p*, generalist proportion) on plant defense investment. **(a)** Graphical illustration of variation in *H* and *p*. (**b**) **Empirical assumption 1:** specialist herbivores are less impacted by a given plant defense compared with generalist herbivores (Cornel and Hawkins 2003; Ali and Agrawal 2012; see also our experiment data in supplementary materials). (**b1**) Specialist damage (*H_s_*) decrease less than linearly with defense investment (*H_s_* = *H*_0_(1 – *γ^a^*), *a* > 1). (**b2**) Generalist damage (*H_g_*) decrease steeper than *H_s_* (*H_g_ = H*_0_ (1 — *γ^b^*),*b* ≥ 1, *b* < *a*, see supplementary materials for the case where *b* = 1). (**c**) **Empirical assumption 2:** allocation cost (C) increase with defense investment convexly (*C* = *γ^b^, b* > 1) (Fagerstrom 1989; Fornoni et al. 2004). **(d)** Marginal benefit 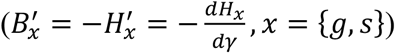 of defense investment (*γ*) increase with increasing *H*_0_. (**d1**) Marginal benefit against specialists is 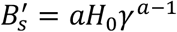. (**d2**) Marginal benefit against specialists is 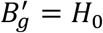. (**e**) Marginal cost of defense could be convex, concave or linear given *b* > 1: *C* = *bγ*^*b*-1^. (**f**) Conditions of interactive effect of herbivory degree (*H*_0_) and generalist proportion (*p*) on optimal defense investment (*γ*\*). If 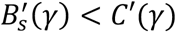 holds true given *γ* satisfies 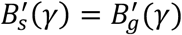, which may occur when herbivory degree (*H*_0_) is relatively low, the optimal defense investment against generalist 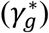 is greater than that of specialist 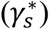. Thus, defense investment should increase with increasing generalist proportion. If 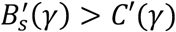 holds true given *γ* satisfies 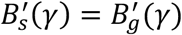, which may occur when herbivory degree (*H*_0_) is relatively high the optimal defense investment against generalist 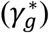 is smaller than that of specialist 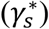. Thus, defense investment should decrease with increasing generalist proportion.

Here, we test the hypothesis that herbivory degree and the ratio of generalist to specialist herbivores interact on the evolution of plant defense (Fig. 1). Specifically, we predict that with increasing proportion of generalist herbivores plant defense should increase when herbivory degree is low; while decrease when herbivory degree is high. To do so, we first use a simple model to formalize the notion of the interaction between herbivory degree and the ratio of generalist to specialist herbivores on plant defense and derive a metric essential for quantifying the relative levels of herbivory. We then explore and illustrate a series of scenarios to predict the pattern of optimal plant defense as a result of interactive effect of change in both herbivory degree and generalist proportion. Lastly, we show how our model relates to a much larger body of works on the geographical pattern of plant defense and herbivory and them can be put into our general theoretical framework.

## Methods

We proposed a modified model for the evolution of plant defense against generalist and specialist herbivores extended from a classic cost-benefit model of plant defense (Lankau, 2007). The general form of this model is as follows:

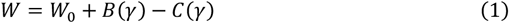

where *W* is the fitness of plant, *γ* is the level of plant defense investment, *W*_0_ is the fitness of a non-defended plant genotype with no defense investment, *B*(*γ*) is the increase in plant fitness related to benefits of defense investment, and *C*(*γ*) is the decrease in plant fitness due to cost of defense investment. It is assumed that plant gains benefit from defense by reducing the herbivory degree it is suffering, and thus *B*(*γ*) is substituted by -*H*(*γ*), in which *H*(*γ*) represents for the realized herbivory damage given certain level of plant defense investment:

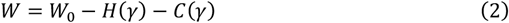

Plant defense investment *γ* in this model is assumed to be normalized, so its value ranges from 0 ≤ *γ* ≤ 1.

We then make further assumptions regarding the characteristic of herbivory and cost functions of plant defense investment.

### Assumptions and empirical background

**Firstly**, generalist herbivores are assumed to be more affected by given levels of plant defense than specialist herbivores (Bergelson *et al.,* 2001; Cornell & Hawkins, 2003; Ali & Agrawal, 2012). This assumption is modeled as relative convexity and slope of herbivory curves of generalist and specialist as below (i.e. magnitude of shape and slope parameters in Eq.3 and Eq.4).

The herbivory damage by generalist herbivore is formulated as:

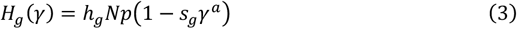

The *h_g_Np* term describes herbivory damage by generalists in absence of plant defense, where *h_g_* is the herbivory damage by a single generalist to a non-defended plant genotype, *p* is the generalist proportion among herbivores, and *Np* represents for population density of generalist herbivores. The (1 – *s_g_γ^a^*) term describes feeding rate of generalist herbivores as a decreasing function of plant defense investment, in which *s_g_* is a slope parameter and *a* is a shape parameter, both describe how feeding rate decrease per unit defense investment.

The herbivory damage by specialist herbivore is formulated as:

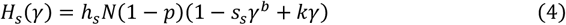

where *h_s_* is the herbivory damage by a single specialist to a non-defended plant genotype, 1 — *p* is the specialist proportion among herbivores, and *N*(1 — *p*) represents for population density of specialist herbivores. In the feeding rate term, *s_s_* and *b* are the slope and shape parameter respectively. The extra term *kγ* describes the increase in specialist feeding rate when plant defense attracts specialist herbivore (e.g. defensive chemicals act as cue for oviposition or feeding simulants). If *k* > 0, herbivory damage by specialists increase with plant defense and peak at intermediate level (shape like a quadratic function), which represents for the case where specialist is attracted by plant defense. If *k* = 0, *H_s_*(*γ*) decrease monotonically, corresponding to the case which specialist are not attracted.

To model the different responses of generalist and specialist to given levels of plant defense (e.g. specialists are more resistant to plant defense than generalists), we assumed that slope and shape parameters should satisfy either *b* > *a* or *s_s_* < *s_g_* or *k* > 0. The condition *b* > *a* implies that herbivory curve of specialist is more convex than generalist; the condition *s_s_* < *s_g_* states that herbivory curve of specialist is flatter than generalist; and the condition *k* > 0 states that specialists are attracted by plant defense while generalists are deterred.

For simplicity, we further assumed the herbivory damage to non-defended plant by a single generalist and specialist are equal, namely *h_g_* = *h_s_* = *h*_0_. Then the total herbivory damage by both generalist and specialist to non-defended plant can be expressed as *H*_0_ = *h*_0_*N* = *h_g_Np* + *h_s_N*(1 — *p*). Here *H*_0_ represents for the maximal fitness loss due to overall herbivory damage, which we named as “ideal herbivory damage”. On the contrary, *H*(*γ*) in Eq. 2 is “realized herbivory damage”. In the later discussion part, we will use ideal herbivory to represent for local gradient of herbivory degree to make predictions rather than using realized herbivory, because realized herbivory itself is coupled with defense level of plant and many other factors. We will also explore the condition where *h_g_* ≠ *h_s_* in the supplementary materials and we will illustrate that major outcomes hold true for such case.

**Secondly**, defense cost is assumed to increase monotonically with defense investment. This assumption is represented by the following relationships between costs and defense investment:

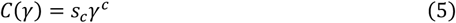

where *s_c_* is a slope parameter and it also denotes the maximal allocation cost of a plant as *C*(1) = *s_c_. c* is the shape parameter of cost function. Some empirical evidence regarding the shape of cost functions has lead previous models to assume that cost function is linear (Bergelson *et al*., 2001). However, resent study have revealed that nonlinear cost functions may be more common than previously expected. For instance, a previous study showed that cost function of defense (i.e., estimated as reduced growth) can vary from being almost linear to being a concave upward function of defense investment (Skogsmyr & Fagerström, 1992). In our model, we examine the cases of linear cost function (*c* = 1), convex cost function (0 < *c* < 1) and concave cost function (*c* > 1).

In summary, the complete expression of the model is as follows:

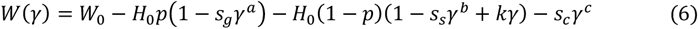

### Detecting the interaction between herbivory degree and generalist proportion

The goal of model evaluation is to find out the conditions where interactive effect on optimal defense level exist between herbivory degree and the ratio of generalist to specialist. To do so, we numerically resolved for optimal defense level 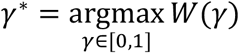 and partial derivatives of optimal defense 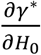 and 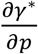. If the interactive effect exists, we expect 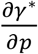 (or 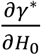) to have different sign given high and low *H*_0_ (or *p*). On the contrary, if there is no interactive effect, 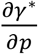 (or 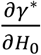) should always be positive or negative regardless of *H*_0_ (or *p*) value.

Technically, optimal defense level *γ*\* was calculated by evaluating 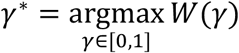 given *γ* in a predefined discrete range (see Table S1). And the partial derivatives 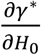 and 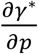 were approximated by discrete expressions 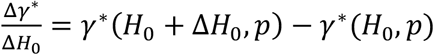 and 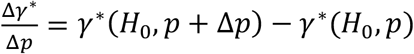, where Δ*H*_0_ and Δ*p* are simulation intervals (see Table S1). We repeated the above calculations for each set of parameters in orthogonal combinations of all parameter values in their predefined range (Table S1), so that we can find the conditions where interactive effect on optimal defense level exist.

## Results

We found the effect of generalist proportion on the optimal levels of plant defense depends on herbivory degree (Fig. 2). Specifically, herbivory degree is divided by a threshold (formulated as *H_T_*, 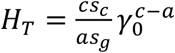 where *γ*_0_ is the solution to equation 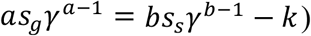 into low and high. When herbivory degree is high (*H*_0_ ≥ *H_T_*), increase in generalist proportion leads to decrease in optimal defense level (Δ*γ*\*/Δ*p* ≤ 0); and when herbivory degree is low (*H*_0_ < *H_T_*), increase in generalist proportion leads to increase in optimal defense level (Δ*γ*\*/Δ*p* > 0). The above result indicates that herbivory degree and the ratio of generalist to specialist do have an interactive effect on the optimal defense level.

**Figure 2.**
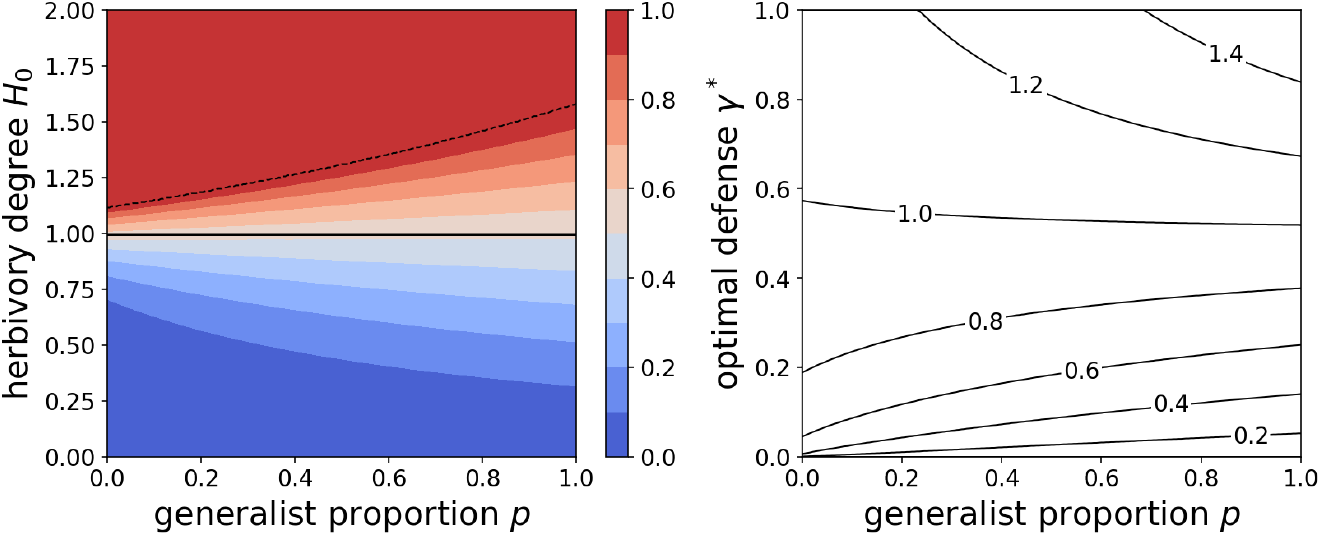
Interactive effect of herbivory degree and generalist proportion on the optimal levels of plant defense. **Left**, contour plot of optimal defense levels. Color indicates the optimal defense level *γ*\*. Solid line is the threshold dividing herbivory degree into low (below the line) and high (above the line). Dashed line indicates that defense level reaches its maximal value (*γ*\* = 1) beyond this line. **Right**, cross sections of contour plot on the left at specific *H*_o_ values. Numbers on the lines indicate the corresponding *H*_o_ values. This pair of plots corresponds to the following parameter setting: *a* = 1.2, *b* = 1.7, *c* = 1.7, *s_g_* = 1, *s_s_* = 1, *k* = 0, *C_max_* = 1.

We found concave cost function (*c* > 1, marginal defense cost increases as defense level increases) is necessary for the existence of interaction (Fig. S1). And in the case of concave cost function, *bs_s_ – as_g_* >*k* is a sufficient condition for the interactive effect to exist. The above expression states that in case *k* = 0 (i.e. plant defense deters specialists), whenever specialists are more resistant to plant defense than generalists, there should be an interactive effect between herbivory degree and generalist proportion; and in case *k* > 0 (i.e. plant defense attracts specialists), it states that attraction effect should be no more than the difference in resistance to plant defense between specialist and generalist for the interactive effect to exists.

## Discussion

Although herbivory degree and generalist-to-specialist ratio have long been seen as important factors driving defense evolution in plants, they were typically treated as independent by presumption, which might have led to great controversy in related studies. Our results provide the first theoretical support for the interactive effect of herbivory degree and generalist proportion on evolution of plant defense, which integrate many previous studies (Table 1) and put them into a general theoretical context (Fig. 3). In the following discussion, we demonstrate how our interactive framework may help empiricists to resolve debates in two specific areas: latitudinal patterns of plant defense and defense evolution of invasive plants.

**Figure 3.**
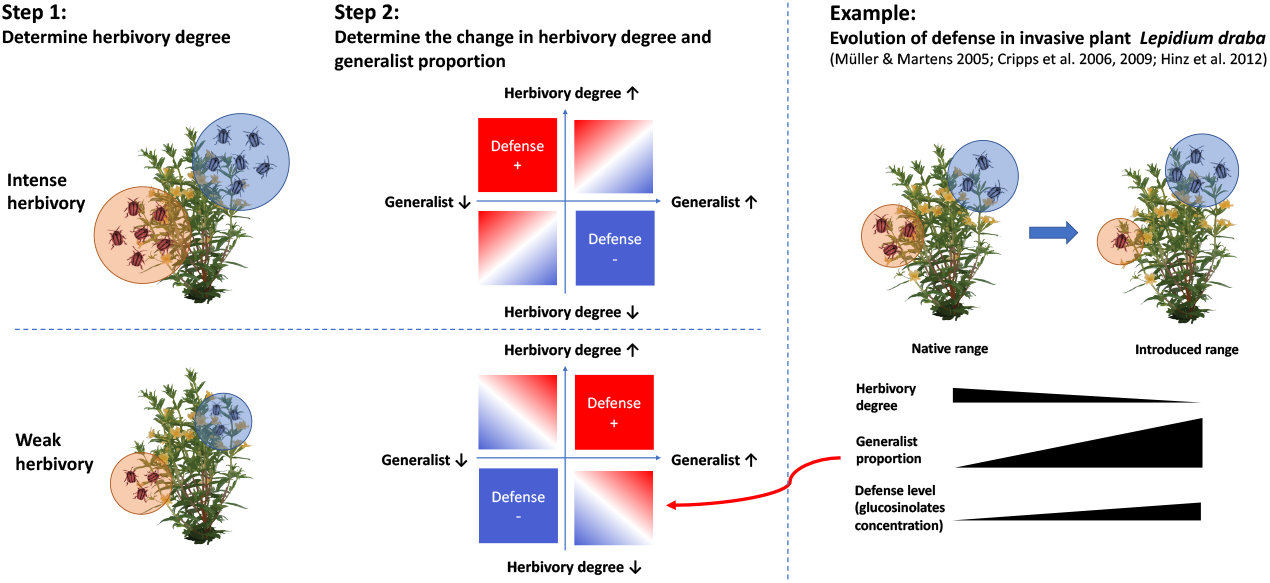
Practical guide to applying interactive effect into predicting the plant defense evolution. In step one, herbivory degree is divided into weak or intense according to herbivory threshold shown in figure 2. In step two, direction and strength of selection gradient on defense is predicted using the change in herbivory degree and generalist proportion. Red color in quadrat plot indicates plant evolves higher level of defense, and blue color indicates lower level of plant defense. A mixture of red and blue color implies that plant may evolve either higher or lower defense according to relative importance of herbivory degree and generalist proportion (see Example on the right)

**Table 1.** Examples of applications and cases supporting our interactive model.

| Scenario | Herbivory degree | Change in herbivory degree | Change in generalist proportion | Predicted change in defense level | Previous theories | Cases |
| --- | --- | --- | --- | --- | --- | --- |
| Plant invasions<sup>1</sup> | High | Decrease | Increase | Decrease | EICA | *Hypericum perforatum* (Maron *et al.*, 2004; Vilà *et al.*, 2005) |
|  | Low | Decrease | Increase sharply | Increase | SDH | *Lepidium draba* (Müller & Martens, 2005; Cripps *et al.*, 2006, 2009; Hinz *et al.*, 2012) |
|  |  | Decrease sharply | Increase | Decrease | EICA | *Silene latifolia* (Wolfe, 2002; Wolfe *et al.*, 2004; Blair & Wolfe, 2004; Elzinga & Bernasconi, 2009) |
| Latitudinal patterns<sup>2</sup> | High | Decrease | Decrease sharply | Increase | - | *Oenothera biennis* (Anstett *et al.*, 2014) |
|  |  | Decrease sharply | Decrease | Decrease | LHDH | *Atriplex spp.*, *Juncus spp.*, *Limonium spp.*, *Spartina spp.* (Pennings *et al.*, 2007) |
|  | Low | Decrease | Decrease | Decrease | LHDH | Cross species (Coley & Aide, 1991); *Acacia falcata* (Andrew & Hughes, 2005)(Coley & Aide, 1991) |
|  | From high to low, crossing threshold | Decrease | Decrease sharply | Increase then decrease, unimodal pattern | - | *Alternanthera philoxeroides* (our unpublished data) |

<sup>1</sup> Here we assume the change in herbivory degree and generalist proportion to be the change from native to introduced range following invasion.
<sup>2</sup> Similarly, we assume the described changes and patterns to be from lower latitude to higher latitude.

### Implication for latitudinal patterns of plant defense

Latitudinal pattern of plant-herbivore interaction is one of the most fascinating but unresolved issues in biogeography. The most studied latitudinal herbivory-defense hypothesis (LHDH) predicted decreased defense towards higher latitude solely based on reduced herbivory degree (Coley & Aide 1991). We point out that ignoring the interactive effect of herbivory degree and generalist proportion could lead to confusion, so that many studies in the recent decade have provided conflicting results (Moles *et al.* 2011a, b; Anstett *et al.* 2016).

For example, a recent case study reported increased herbivory degree at higher latitude, and in the meanwhile concentration of major defensive substances declined (Anstett *et al.,* 2015), which conflicts with the prediction of the LHDH. While we found in this case, slope of latitude-versus-herbivore-abundance curve was greater for specialist than generalist, indicating a shift towards lower generalist proportion at higher latitude, and this is supported by a related study of the same author (Anstett *et al.,* 2014). Given the herbivory degree in this case kept in relatively low level, lower generalist proportion in high latitude should select for lower defense.

Moreover, our interactive framework is able to capture almost all possible latitudinal patterns of plant defense that have been reported to date (Moles *et al.* 2011a; Anstett *et al.* 2015, and also see Fig. S2). Thus, we propose that combining herbivory degree and generalist proportion in future studies can be a good start point to merge divergent patterns into an unified framework. A growing number of studies consistently show that diet breath of herbivore insects shift globally towards a higher frequency of specialist insects at lower latitude (Novotny *et al.,* 2002; Dyer *et al.,* 2007; Forister *et al.,* 2015). Current and coming evidences on gradients of generalist-to-specialist ratio should push future studies to pay more attention on combining the herbivory degree and generalist-to-specialist ratio when studying latitudinal herbivory-defense patterns.

### Implication for defense evolution in invasive plants

Herbivory degree and the ratio of generalist to specialist have also been seen as important factors in shaping the evolution of defense in invasive plants. However, many previous theories treat them as separate factors and thus making conflicting predictions. For example, the evolution of increased competitive ability (EICA) hypothesis predicted reduce in defense level based on less herbivore pressure in plants’ introduced ranges (Blossey & Nötzold 1995). While the shifting defense hypothesis (SDH) predicted increase in some specific defense traits against generalist in introduced range due to increased generalist proportion (Müller-Schärer *et al.* 2004; Joshi & Vrieling 2005).

Our interactive model reveals the conditions where the EICA hypothesis or SDH can be applied and thus integrates them into an unified framework (Table 1): if herbivory degree in native and introduced range are both weak, then the EICA fits in the cases where decrease in herbivory degree is sharper then increase in generalist proportion and the SDH fits in complementary cases where increase in generalist proportion outweighs the decrease in herbivory degree (see the example shown in Fig. 3). If herbivory degree are both high, then decrease in herbivory degree and increase in generalist proportion in combination will consistently select for reduced defense level as predicted by the EICA hypothesis and the SDH could not fit into the high herbivory degree scenario. These conditions are consistent with general conclusions of a recent meta-analysis (Zhang *et al.* 2018).

### Future Directions

The interaction between herbivory degree and generalist proportion is largely unexpected by previous studies, but we suspect it can be ubiquitous. Theoretically, such interaction between herbivory degree and generalist proportion roots in the nonlinearity of the marginal benefits of plant defense. As shown in our conceptual diagram (Fig. 1), the marginal benefit curve of generalist intersects that curve of specialist, meaning that defending against generalists is more efficient than defending against specialists given high level of defense, while less efficient given low level of defense. As defense level covaries with herbivory degree, it is reasonable to expect the existence of interaction between herbivory degree and generalist proportion when defense benefits are nonlinear and unidentical. Many studies support the nonlinear and unidentical assumption (Bergelson *et al.,* 2001; Lankau, 2007), so we suspect that such interaction could be prevalent. For example, a review paper by Bergelson et al. (2001) enumerated eight examples of damage-by-herbivore-versus-plant-resistant plots, and all this cases support nonlinearity in benefit of plant defense against herbivores.

Empirically, no controlled experiment has directly detected the predicted interaction between herbivory degree and generalist-to-specialist ratio, but some field observations provide indirect evidence for evidence for the interaction (see cases in Table 1). We call for more attention on finely designed experiments that manipulating gradients of herbivory degree and the ratio of generalist to specialist simultaneously to test for the interactive effect. Using well documented observations in the field can be an alternative way of testing, and for example invasive plants or widely distributed species across latitudes can be suitable materials.

We note that our study still has some limitations, which will direct our future works. We applied an optimization approach, which features in its simplicity and is usually sufficient to model systems at equilibrium. However, optimization models may be less predictive in a dynamic time scale. Future works will focus on understanding the evolutionary dynamics of defense in the light of the ecological feedbacks that are intrinsic to the interaction of plant and herbivores (Lankau, 2007).

## Conclusion

Our model of the interaction between herbivory degree and generalist proportion suggests that considering these factors simultaneously can improve predictions of plant defense levels. This model provides a general theoretical framework for analysis of genetically based intraspecific variation of plant defense level, helps to explain previous experimental results, and has important implications for the biogeography of plant defense and the evolution of defense in invasive plants.

## Acknowledgements

This work is supported by Hui-Chun Chin and Tsung-Dao Lee Chinese Undergraduate Research Endowment (grant No. 17929 to YP), and National Natural Science Foundation of China (grant No. 41771053 to XP). We thank Shurong Zhou, Ingrid Parker, Dana Blumenthal, Mu Liu and Xiang Liu for their comments and advices on early draft of this manuscript.

## Author contributions

Y.P. and X.P. conceived the ideas. Y.P. conducted modeling and simulation. Y.P. wrote the paper with major inputs from X.P., L.D.B.F. and B.L..

## Code availability

All codes and scripts in this work are available on GitHub repository (https://github.com/Augustpan/Defense_Evo_Model).

## Notes

### Competing Interest Statement

The authors have declared no competing interest.

