## Supplementary materials for "Interactive effect matters: a combination of herbivory degree and the ratio of generalist to specialist better predicts evolution of plant defense"

### Supporting information

### Figure S1

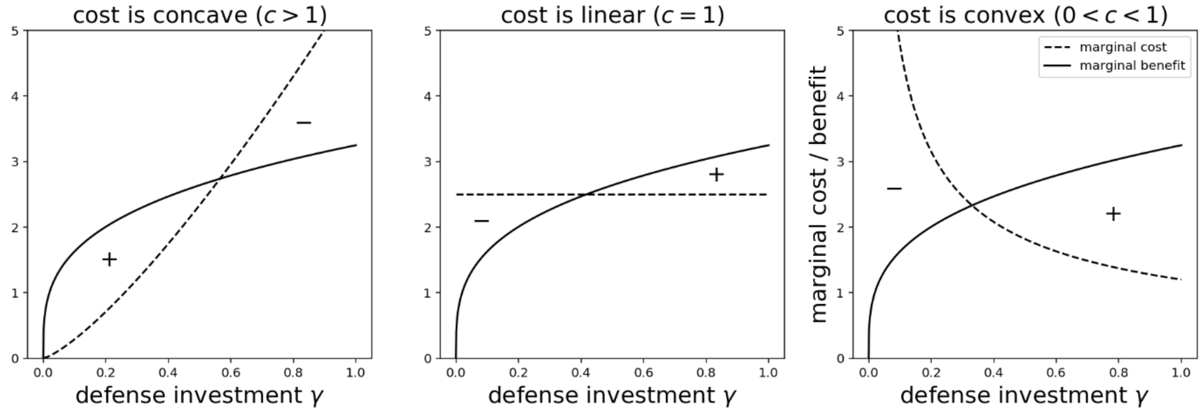


**Figure S1.** Cost-benefit analysis in cases of concave, linear and convex cost functions. In case of linear or convex cost function, defense investment minimized at intermediate level $\gamma \in (0,1)$, and therefore optimal defense level should be either 0 or 1. So there could not be interaction between herbivory degree and the ratio of generalist to specialist given $0 < c \leq 1$.

### Figure S2

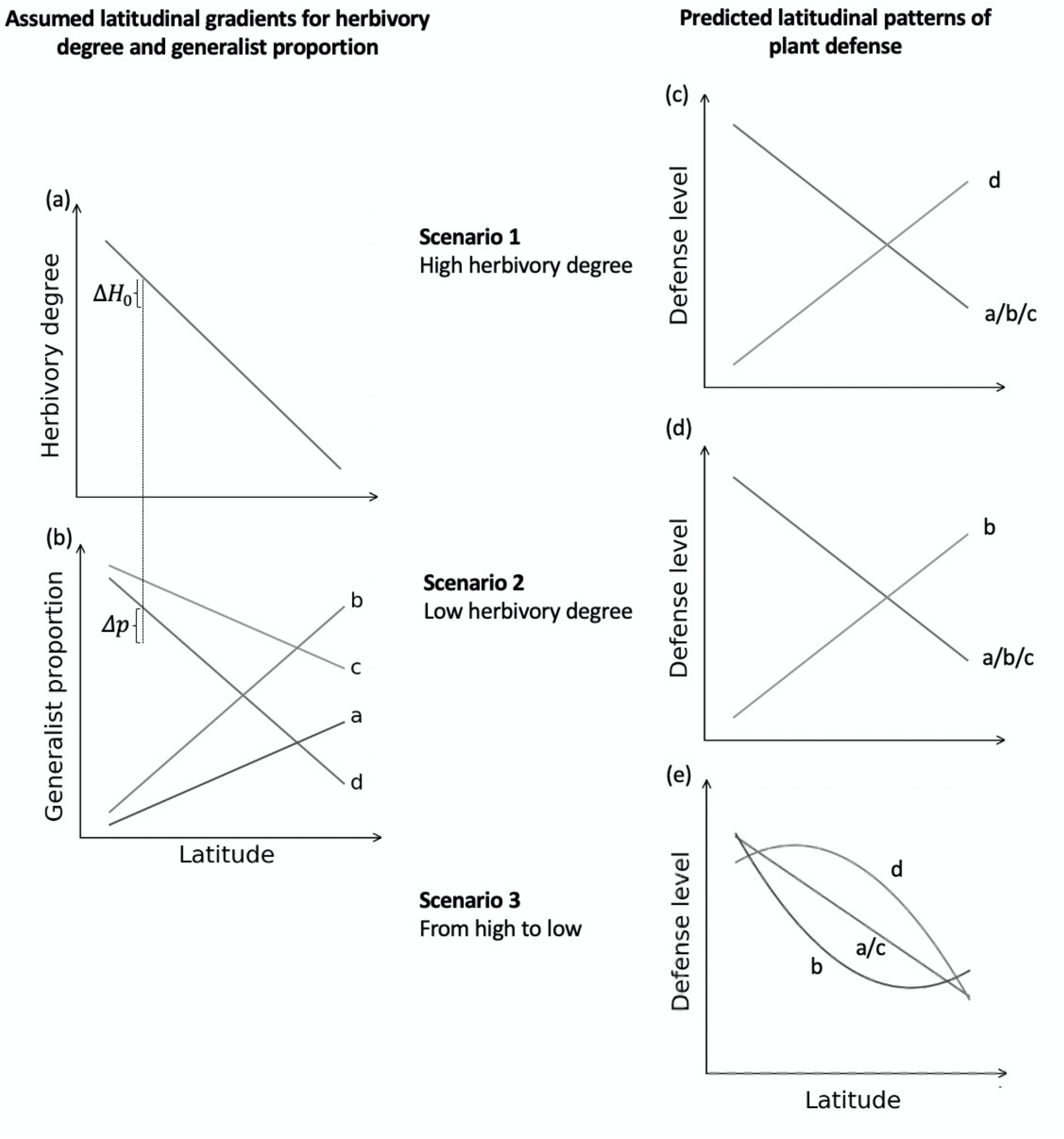


**Figure S2.** Predicted latitudinal patterns of plant defense with our interactive model. (**a, b**) assumed latitudinal gradients for herbivory degree and generalist proportion. (**c, d, e**) predicted latitudinal patterns of plant defense. The label nearby each curve indicates the corresponding pattern is generated with assumed gradient herbivory degree in combination with the gradient of generalist proportion with the same marker.

**Table S1**. Parameters definition and simulation setup

| Notation | Definition | Simulation setup |
|---|---|---|
| $\gamma$ | Normalized plant defense investment | $\gamma \in [0,1]$ at 0.001 interval |
| $N$ | Sum of population size (density) of all herbivores, $h_0 N = H_0$ | / |
| $p$ | Generalist proportion | $p \in [0,1]$ at 0.01 interval |
| $H_0$ | Herbivory damage to non-defended plant by both generalist and specialist | $H_0 \in [0,2]$ at 0.01 interval |
| $h_0$ | We assumed $h_g = h_s = h_0$ | / |
| $h_g$ | Herbivory damage to non-defended plant by a single generalist herbivore | / |
| $h_s$ | Herbivory damage to non-defended plant by a single specialist herbivore | / |
| $s_g$ | Slope parameter of herbivory damage curve for generalists | $s_g \in (0,1]$ at 0.1 interval |
| $s_s$ | Slope parameter of herbivory damage curve for specialists | $s_s \in (0,1]$ at 0.1 interval |
| $s_c$ | Slope parameter for cost function | Fixed at $s_c = 1$ |
| $W_0$ | Fitness of a non-defended plant genotype without herbivory | Fixed at $W_0 = 0$ |
| $a$ | Shape parameter for generalist herbivory function | $a \in (1,3]$ at 0.1 interval |
| $b$ | Shape parameter for specialist herbivory function | $b \in (1,3]$ at 0.1 interval |
| $c$ | Shape parameter for cost function | $c \in (0,3]$ at 0.1 interval |
| $k$ | Attraction parameter, describe how specialists are attracted by plant defense | $k \in [0,1]$ at 0.1 interval |

**Table S1. Comments: justifications for ranges of parameters (simulation setups)**

We set $H_0 \in [0,2]$ because in most combinations of parameters, optimal defense level reach its maximal values $\gamma^* = 1$ when $H_0 = 2$. And in those cases where optimal level does not maximize at $H_0 = 2$, we further increase the $H_0$ value until it reaches maximum.

$s_g \in (0,1]$ and $s_s \in (0,1]$ is the definition space for the two parameters, so such simulation setting captures full range of all possible values.

We fix $s_c = 1$ because changing the value of this parameter only scales Eq. 6 by a constant and will surely not change any major outcomes. And the reason for fixing $W_0 = 0$ is the same.

The range of shape parameters $a, b, c$ is chosen according to empirical data from Bergelson et al. 2001 and Lankau 2007. We extracted data from figures, and fit the relation between defense level and herbivory damage as well as the relation between defense level and plant fitness to get estimations of parameter $a, b, c$. We set the upper bound of the numerical simulation range to be more than 150% of the biggest fitted values to include most of the possible cases in real world.

We set $k \in [0,1]$ because when $k > 1$, such case is so extreme that plant defense will not having any benefit on defending against specialists even if the defense level is maximized.
